# Progression of *ampC* amplification during de novo amoxicillin resistance development in *E. coli*

**DOI:** 10.1101/2024.05.24.595737

**Authors:** Luyuan Nong, Martijs Jonker, Wim de Leeuw, Meike T. Wortel, Benno ter Kuile

## Abstract

Beta-lactam antibiotics are the most applied antimicrobials in human and veterinarian health care. Hence, beta-lactam resistance is a major health problem. Gene amplification of AmpC beta-lactamase is a main contributor to de novo β-lactam resistance in *E. coli*. However, the time course of amplification and the accompanying DNA mutations are unclear. Here, we study the progression of *ampC* amplification and *ampC* promoter mutations in the evolution of resistance by stepwise increasing amoxicillin concentration. *AmpC* promoter mutations occur by day two, while the amplification by a factor of approximately eight occurs after more than six days of amoxicillin exposure. The combination of amplification and promoter mutations increase *ampC* mRNA level by an average factor of 200 after 22 days. An IS1 insertion was identified in the amplification junction, suggesting the amplification is facilitated by mobile genetic elements transposition. In order to identify the essential genes for *ampC* amplification, the chromosomal mutations of strains with induced amoxicillin resistance were compared a similarly evolved resistant Δ*ampC* knockout. The evolved Δ*ampC* contained several resistance mutations that were absent in the WT, which accumulated more mutations in stress response genes. The amoxicillin evolved Δ*ampC* does not show amplification of the fragment around the original *ampC* position but exhibits a large duplication or triplication at another position, suggesting selection of genes to amplify is essential for resistance adaption.

**IMPORTANCE:** Amoxicillin is one of the most used antimicrobial against bacterial infections. DNA fragments containing *ampC* are amplified upon prolonged and stepwise increasing exposure to amoxicillin. These *ampC* amplification fragments have been identified in extended-spectrum beta-lactamases (ESBLs) plasmids, which are considered the main cause of beta-lactam resistance. Understanding the progression of *ampC* amplification enables amoxicillin resistance prevention. In this study, we show the time course of two important factors for *ampC* transcription enhancement, *ampC* amplification and *ampC* promoter mutations, during de novo amoxicillin resistance evolution. We propose that the transposon IS1 contributes to the amplification and that the sigma factor 70 regulates *ampC* overexpression.

## INTRODUCTION

Although alternative antibiotics are available (1)(2), beta-lactam antimicrobials remain the most commonly used antibiotics for human infection therapy (3). Exposure to sublethal levels of antibiotics is the main reason for resistance emergence (4). When the effectiveness of antibiotics is reduced, exposure to sublethal levels during antibiotic treatment is unavoidable (5). The risk of non-lethal concentrations occurs especially in the veterinary sector when livestock is treated with antibiotics added to water or feed. Under long-term exposure, antimicrobial resistance may accumulate in the microbiome and this resistance can be transmitted to human pathogens through horizontal gene transfer.

AmpC beta-lactamases (AmpC) are widely distributed cephalosporinases causing beta-lactam degradation (6). The *ampC* gene can be encoded by both chromosomal and plasmid DNA. Unlike extended-spectrum beta-lactamases (ESBLs) which are considered the main cause of the prevalence of beta-lactam resistance (7), the overexpression of *ampC* is underestimated. In *Escherichia coli*, the chromosomal *ampC* gene is poorly expressed in the wildtype, as it lacks the transcription activator AmpR (8). However, after laboratory evolution of amoxicillin resistance, the transcription of *ampC* is enhanced more than 100-fold compared to the naive strain, mainly because of two reasons: *ampC* promoter mutations and gene multiplication (9). The *AmpC* gene is controlled by a weak promoter (P*_ampC_*) which has three main elements, a - 10 box, a -35 box and an attenuator (10). The mutations C > T in the -10 box at position -11 and T > A in the -35 box at -32 are conservative, increasing AmpC production by 21- and 7- fold, respectively (10). In addition, *ampC* gene amplification plays a considerable role in its transcriptional enhancement (11). Chromosomal *ampC* amplification was first reported increasing resistance (12). This amplification is a RecA-independent event occurring in tandem (13)(14). Duplicated antibiotic resistance genes could undergo horizontal gene transfer in microbial communities, with mobile genetic elements serving as a vehicle (15). A chromosomal DNA fragment containing the *ampC* gene was amplified from strains made resistant to amoxicillin by exposure to step-wise increasing concentrations and isolated with plasmid isolation techniques (16). This fragment can be exchanged between *E. coli* strains by horizontal gene transfer. A similar *E. coli* chromosomal fragment harboring *ampC* and two nearby genes can be identified in several ESBL plasmids isolated in broiler production (17). This suggests that *ampC* amplification may occur in *E. coli* developing beta-lactam resistance and that subsequently the fragment is incorporated in ESBL plasmids.

In addition to inactivation of the antibiotic through AmpC, various other molecular mechanisms can contribute to beta-lactam resistance, including active efflux pumps, decreased influx, and target site modification (18). Multiple point mutations were demonstrated to directly contribute to resistance, such as mutations in the efflux pump AcrAB-TolC (19) and the outer membrane porin OmpC/OmpF (20). Besides, some mutations that alter metabolism also confer to resistance (21)(22). For example, a mutation in the 2- oxoglutarate dehydrogenase (*sucA*) gene raised carbenicillin resistance through lower basal respiration, thereby avoiding metabolic toxicity and reducing lethality.

To understand the competitive, synergistic and epistatic effects of DNA mutations associated with *ampC* amplification, this study investigates the time course of mutations in the chromosome and *ampC* amplification. Using the de novo development of amoxicillin resistance in *E. coli* as a model, this study addresses five questions: 1) What is the time course and pattern of *ampC* amplification? 2) Is the amplified *ampC* fragment always the same, or can the length vary? 3) Is the *ampC* copy number the primary factor determining the AmpC activity? 4) If the *ampC* gene is removed, does the amplification of the fragment around it still occur? 5) Is the pattern of mutations accompanying development of resistance different in the presence and absence of *ampC*? Answering these questions provides insights into the complex dynamics of beta-lactam resistance development and the role of *ampC* amplification in this process.

## RESULTS

### Relationship between AmpC activity and amoxicillin resistance

To investigate the role of AmpC in amoxicillin resistance, nine replicates of wild type (WT) and six of *ampC* knock-out mutants (Δ*ampC*) were made highly resistant by growing them at stepwise increasing sublethal amoxicillin concentrations (Fig.1a). The acquisition of resistance by Δ*ampC* occurs significant slower than by the WT. Correspondingly, the minimal inhibitory concentration (MIC) reached at least 1024 mg/L in WT and only approximately 128 mg/L in Δ*ampC* (Fig.1b). The AmpC activity was determined using the chromogenic substrate nitrocefin in the presence of intact live cells, as the enzyme functions as an ectoenzyme (9). Compared to the activity of the naive wildtype, the activities encountered in the evolved WT increased by a factor of 39-156 in the final incubations (Fig.1c). The log_2_ transformed fold change of MIC and AmpC activity exhibited a linear relationship with R^2^=0.730 (Fig.1d). This indicates that de novo amoxicillin resistance in *E. coli* can be largely attributed to the increase in AmpC activity.

### *AmpC* gene amplification during the evolution of resistance

The *ampC* gene, coding for a beta-lactamase, can be amplified when the cells are exposed to stepwise increasing non-lethal levels of amoxicillin (16). In order to determine the moment that this amplification takes place, the copy number of chromosomal *ampC* was measured at three-day intervals using qPCR in all evolved WT replicates with naive WT as reference (Fig.2a). The first amplifications were observed on day nine. In the first six days, the *ampC* copy number did not increase in any of the replicates (Fig.2b). After that, in only one replicate did the copy number increased gradually, in all other replicates an initial jump of a factor eight or more was observed. The median time point for amplification was around day 12, the last single copies were seen on day 16, and at days 19 and 22 all copy numbers ranged from 8 to 19.

In order to determine the size and composition of the DNA fragment containing *ampC* that is amplified, whole genome sequencing was performed on all replicates of WT and three replicates of Δ*ampC* at day 22. Seven different fragments were found, of which five were unique and two fragments were observed twice (Fig.3a). The length of the amplified region ranged from 7.5 kb to 13.4 kb. Five fragments shared the same left terminal, located within the *ecnB* promoter or reading frame, indicating that a preferred target exists for initiation of the amplification. One termination sequence is shared by 3 fragments, each with a different initiation site. All amplified fragments contain the *sugE*, *blc* and *ampC* genes. *SugE* codes for efflux transporters in the multidrug resistance family (23), while *blc* codes for an outer membrane lipoprotein (24). Both *sugE* and *blc* may contribute to amoxicillin resistance.

We tested whether the amplification fragments are connected to each other by designing the forward primer to bind to the start and the reverse primer to bind to the end sequences of the amplified fragments. The junction between two copies of amplification contigs was cloned by PCR and confirmed by Sangar sequencing. If the product shows a connection, either the fragment has become circular, or two or more fragments are connected. Two types of junctions were found (Fig.3b). In four of nine replicates, two amplification contigs directly connect, with 0 to 7 bp overlapping base pairs. In the other five, an IS1 transposon of 768 bp, was inserted in the junctions with also 0 to 7 bp homology base pair between one amplification fragment and the IS1 element. Interestingly, all of replicates with IS1 inserted into amplification junction have one of their flank sites located in the *ecnB* promoter or reading frame.

In evolved Δ*ampC*, there was no amplification near the original *ampC* position, indicating that the other genes in the amplification fragment of evolved WT contribute much less or not at all to amoxicillin resistance (Fig. 3c). Instead, a large chromosomal duplication or triplication occured in a similar position, between base pairs 387,800 and 1,391,170 on the chromosomal map. Genes in this region that may contribute to amoxicillin resistance include *acrA*, *acrB*, *ompF*, sulA, *ftsZ*.

**Fig. 1.**
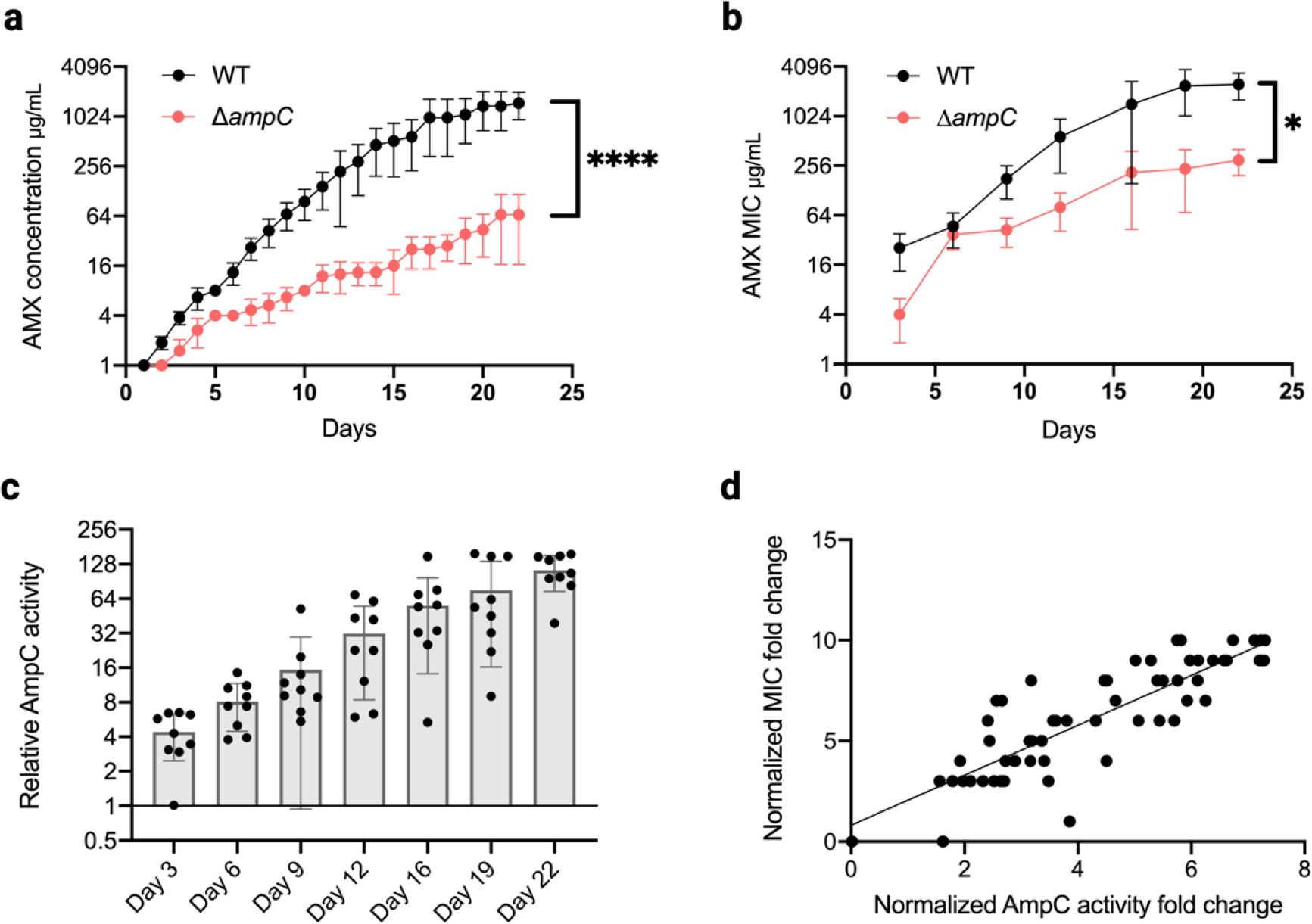
Evolution of amoxicillin resistance. a. Amoxicillin concentration in the cultures. **b** Measurement of MIC for amoxicillin. Data in **a, b** are shown as means ± SD, statistical significance was determined with Wilcoxon signed-rank test, ****p<0.0001, *p<0.05. **c** Measurement of AmpC activity. Each point represents the mean of three technical replicates. **d** Relationship between MIC and AmpC activity. The fold change of MIC and AmpC activity were transformed by log_2_ and fitted with linear regression (R^2^=0.730).

**Fig. 2.**
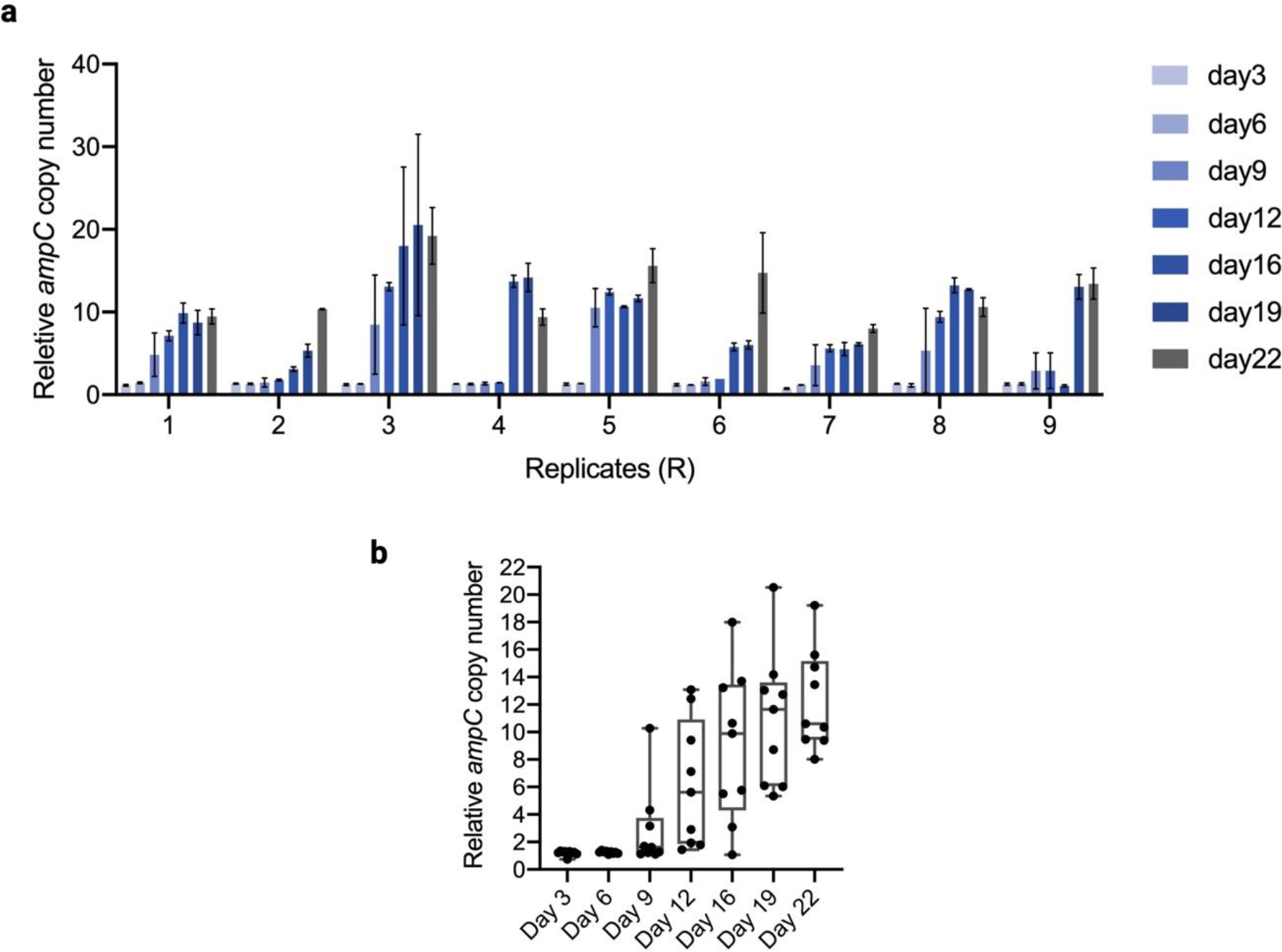
ampC copy number at different time points during the amoxicillin evolution. a. The copy number measurement in each biological replicate. **b** The distribution of *ampC* copy number at each time point in nine biological replicates. Each point is the mean of three technical replicates.

**Fig. 3.**
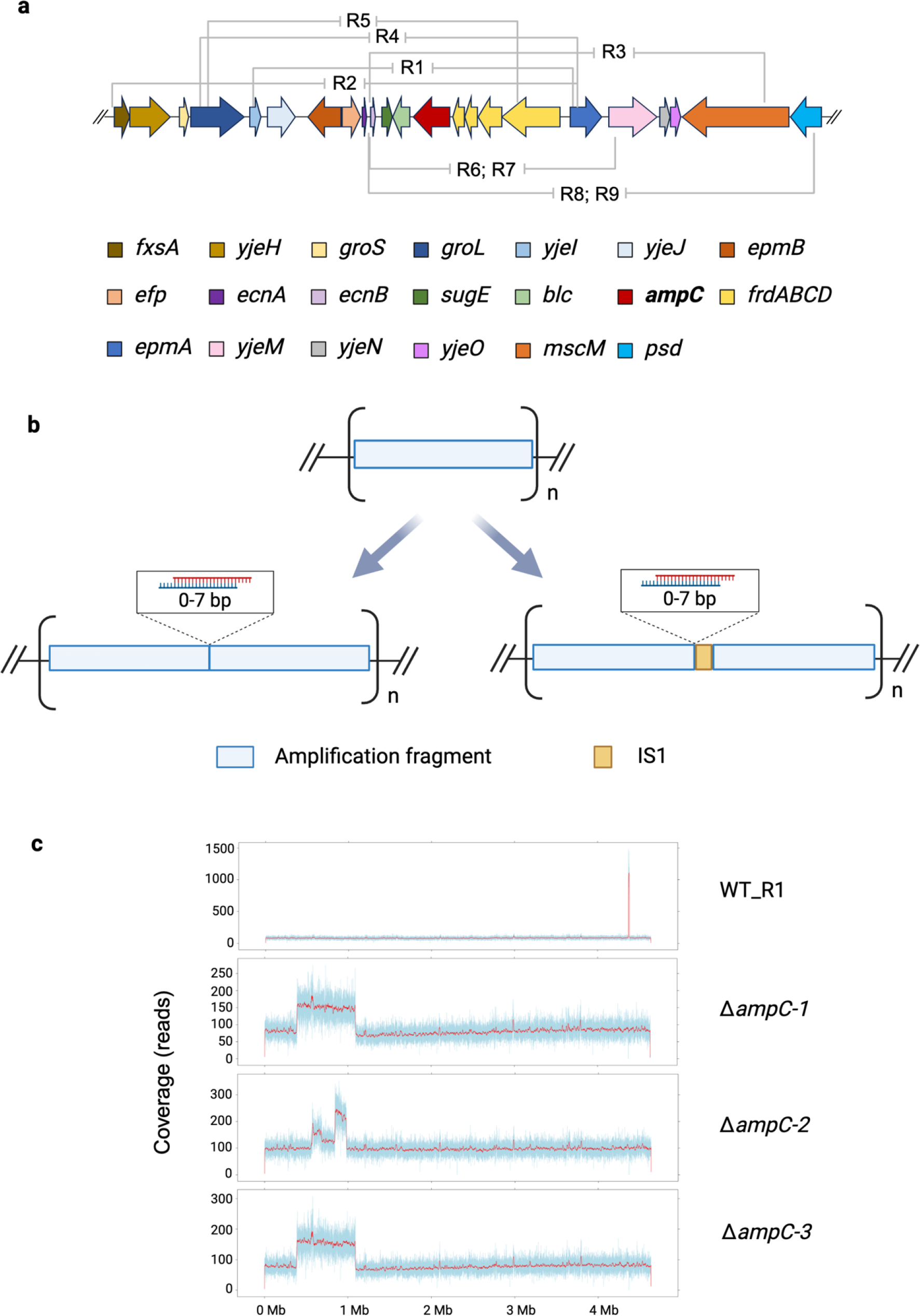
Amplification regions in the evolved strains. a. The various amplification fragments in evolved WT. **b** Junctions between fragments after amplification. Two situations are observed. Four replicates’ amplified fragment connected tandemly with 0-7 base pair overlapping. In five replicates’amplified fragments have an IS1 transposon insertion in their connection junctions with 0-7 overlapping base pair between amplified fragment and IS1. **c** Read counts in whole genome sequencing of evolved Δ*ampC* strains. Large regions with duplication or triplication occurred.

### DNA mutations in the evolved WT and Δ*ampC*

To find which genes are mutated in association with the amplification of *ampC*, whole genome sequencing was performed on DNA isolated at the moment of *ampC* amplification (AT) and the end of the evolution experiment (FT) in WT strains, as well as the Δ*ampC* strains at FT. Mutations were identified through aligning to *E. coli* genome download from NCBI, eliminating those also present in the control and those observed at frequencies below 10%. Except mutations referred to *ampC*, a total of 99 DNA mutations (84 single-nucleotide polymorphisms, 5 deletions and 10 insertions) were identified in all evolved WT (Table S1, S2). Of these, 32 of point mutations were outside of reading frames. In the evolved Δ*ampC* strains we identified 97 DNA mutations with the same analysis workflow (96 single-nucleotide polymorphisms and one deletion), 55 of them are outside of gene reading frames (Table S2, S3). The genes of interest are divided in two functional groups, stress response and cell envelope, and are presented in a heatmap (Fig. 4).

**Fig. 4.**
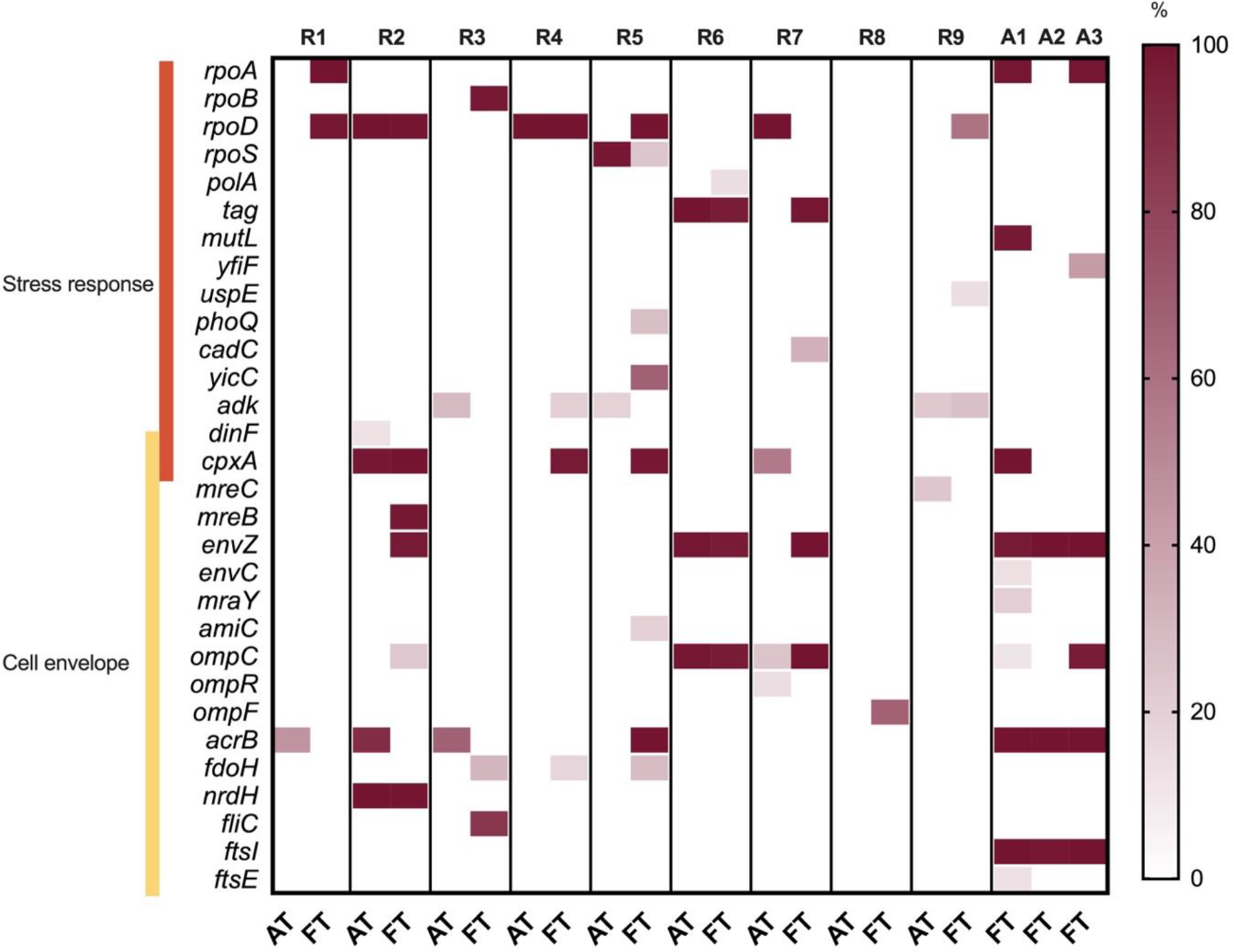
**Mutations associated with *ampC* amplification in amoxicillin evolution**. The color intensity indicates the frequency of mutations in the genes indicated. The time points are the *ampC* duplication (AT) and the end of the experiment inducing resistance (FT). R1 to R9 are evolved replicates of WT, A1 to A3 are evolved replicates of Δ*ampC*.

**Fig. 5.**
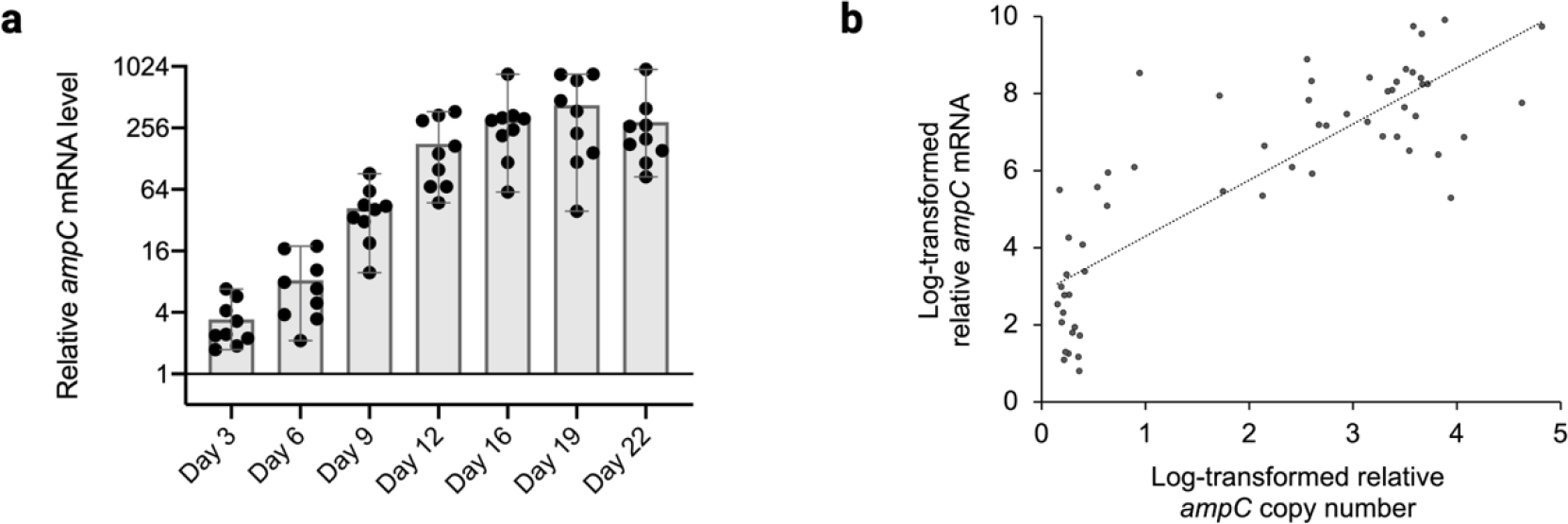
AmpC transcription level in amoxicillin evolution. A. Fold-change in *ampC* expression compared to the native WT. Each point represents the mean of three technical replicates. **b** The relationship between *ampC* copy number and *ampC* mRNA level. The log-transform relative copy number and mRNA were fitted into a linear trendline.

Not one mutation is present in all replicates. In the evolved WT, several mutations associated with stress response are identified coding for sigma factors (*rpoA*, *rpoB*, *rpoD*, and *rpoS*). Most mutations are observed in *rpoD*, that contains mutations in six out of nine replicates. The majority of these mutations show an increasing or sustained frequency from the time point of the first amplification observed to the end of the evolution experiment. In one replicate mutations in *rpoD* appear to outcompete those in *rpoS*, as evidenced by a decreased mutation frequency in *rpoS* and a simultaneous increase in *rpoD*. The DNA repair related gene *polA* was mutated in two replicates. Genes related to cellular response to abiotic stimulus (*uspE* /*phoQ*), stress-induced mutagenesis (*yicC*) and biosynthesis of purine ribonucleotides (*adk*) were mutated occasionally. Four genes (*phoQ* /*adk* /*cpxA* /*envZ*) in the network of phosphorus metabolic processes, which are involved in energy and carbon metabolism, had mutations, suggesting a role of the network in building up amoxicillin resistance. The gene *cpxA* codes for a membrane sensor for envelope stress response (25). Its mutations can increase β-lactam resistance in *E. coli* (26). Different mutations with high and increasing frequencies occur in *cpxA* in several replicates, N346K in R2, Q34P in R4, T16_T17 deletion in R5 and K121I in R7.

The mutations in membrane related genes correspond to different mechanisms for amoxicillin resistance, such as active efflux and decreased influx (18). Because amoxicillin blocks cell wall synthesis by inhibiting transpeptidation, the mutations in genes related to cell division (*mreB* /*mreC* /*amiC*) may counter the effects of long-term amoxicillin exposure. The genes *envZ* and *ompR* code for a two-component system that forms a sensor for outer membrane diffusion pores OmpF and OmpC (27). The change of both *ompC* and *ompF* can alter their substrate specificity and restrict the influx of amoxicillin, causing resistance (28, 29). The mutations consisted of P248L in *envZ* and D33fs in *ompC* .These mutations occurred in the same replicates at the same times. The mutations frequency in *ompC* is higher than *ompF*, although the deletion of *ompC* causes a lower MIC against many beta-lactam antibiotics, while the deletion of *ompF* increases them (20). AcrB is a well-known drug efflux pump that affects *E. coli* amoxicillin resistance (30). The mutation in *acrB* was accumulated in three replicates in AT. However, these mutations were absent at the end of the evolution experiment. Both *fdoH* and *nrdH* are involved in the electron transport chain, hence the mutations in these two genes indicate that the *ampC* amplification occurs in parallel with the fitness alteration.

The evolved Δ*ampC* strains tend to exhibit mostly mutations associated with membrane. All Δ*ampC* replicates contained mutations in *ftsI*/*acrB*/*envZ*. The inhibition of FtsI activity by binding of beta-lactam antibiotics is lethal, as this is an essential cell division protein (31). Although the gain-of-function mutations in *ftsI* is well-known for amoxicillin resistance (32), this mutation did not occur in evolved WT strains. In addition, in evolved Δ*ampC*, the proportion of mutations located in *envZ*/*ompC*/*acrB* is relatively higher than in the WT, suggesting an increasing role of other resistance pathways. The mutation in *envC*, which codes for the efflux pump AcrE, occurred occasionally. Although the *acrEF* operon is not essential in *E. coli* when the *acrAB* operon is expressed, when *acrAB* become inactive, the *acrEF* operon complements the function of *acrAB* under stressful conditions (33). This suggests AcrE has the ability to cause resistance. Besides, mutation accumulation was observed in the genes coding for the cell division protein FtsE and the peptidoglycan lipid biosynthesis enzyme MraY in Δ*ampC,* but not in the WT. In regard to the mutations associated with stress response, the mutations in *rpoD* which have a high frequency in evolved WT were not observed in Δ*ampC*. However, the mutations in sigma factor subunit *rpoA* occurred at a higher frequency in Δ*ampC*. Fewer mutations in stress response genes were observed in evolved Δ*ampC* strains than in the WT. Only mutations in DNA mismatch repair protein gene *mutL* and putative methyltransferase gene *yfiF* were found.

### Relationship between *AmpC* mRNA level and *ampC* copy number

To determine the consequences of the observed *ampC* gene copy number increase, *ampC* mRNA levels were measured at the same time points that the *ampC* copy number was ascertained (Fig.5a). Even though there was no gene amplification within the first 6 days of the evolution experiments, a continuous increase of *ampC* mRNA levels was evident, indicating that other factors also enhance *ampC* transcription ahead of *ampC* amplification. During induction of resistance, *ampC* mRNA levels significantly increased in all replicates, ranging from 85 to 961-fold change compared to the WT.

In order to establish the relationship between *ampC* copy number and mRNA level, the *ampC* mRNA level and copy number fold-change were transformed by log_2_ and fitted in linear function (Fig.5b). The *ampC* mRNA level and copy number have a positive correlation.

However, the linear formula does not fit well. This suggests that other factors, in addition to *ampC* copy number, also have considerable impact on *ampC* gene transcription.

### Trajectory of mutations related to *ampC*

Instigated by the evidence above, we further explored how the other mutations affect *ampC* transcription. Besides gene dosage, promoter activity is another crucial factor influencing gene transcriptional levels. Therefore, the *ampC* promoter region was sequenced at 8 different time points during the evolution experiments (Table S4). *AmpC* promoter mutations were observed as early as day 2, affecting the -10 box of the promoter. As evolution progressed, more mutations emerged. However, not all of them were retained until the final days. The first mutation in the *ampC* promoter occurs earlier than *ampC* amplification. Combining this information with the observation that *ampC* mRNA levels increase by about 10-fold prior to *ampC* gene amplification, suggests that the mutated *ampC* promoter is responsible for the initial increase in *ampC* transcription.

The mutations in the *ampC* promoter occurred in three main elements: the -10 box, the -35 box, and the attenuator. In the nine replicates, the mutations in the -10 box and -35 box were conserved, -11 G>A in -10 box and -32 A>T, but not those in the attenuator. In the attenuator area 5 different mutations were found in different replicates at the end of evolution. Additionally, there were a few mutations in other sites that have not been reported before in the *ampC* promoter. To uncover a possible influence of changes in the *ampC* promoter region on the *ampC* copy number, the trajectory of mutations in the *ampC* promoter region was documented (Fig.6). Mutations in the -10 box already occurred by day 2. However, the frequency of this mutation decreased after day 3, accompanied by an increase in the frequency of mutations in the -35 box. After the *ampC* copy number started to increase, mutations in the -10 box and -35 box did not show systematic changes. Mutations in the attenuator and undescribed area were more unpredictable, with their mutation frequency continuously changing throughout the entire evolution process.

**Fig. 6.**
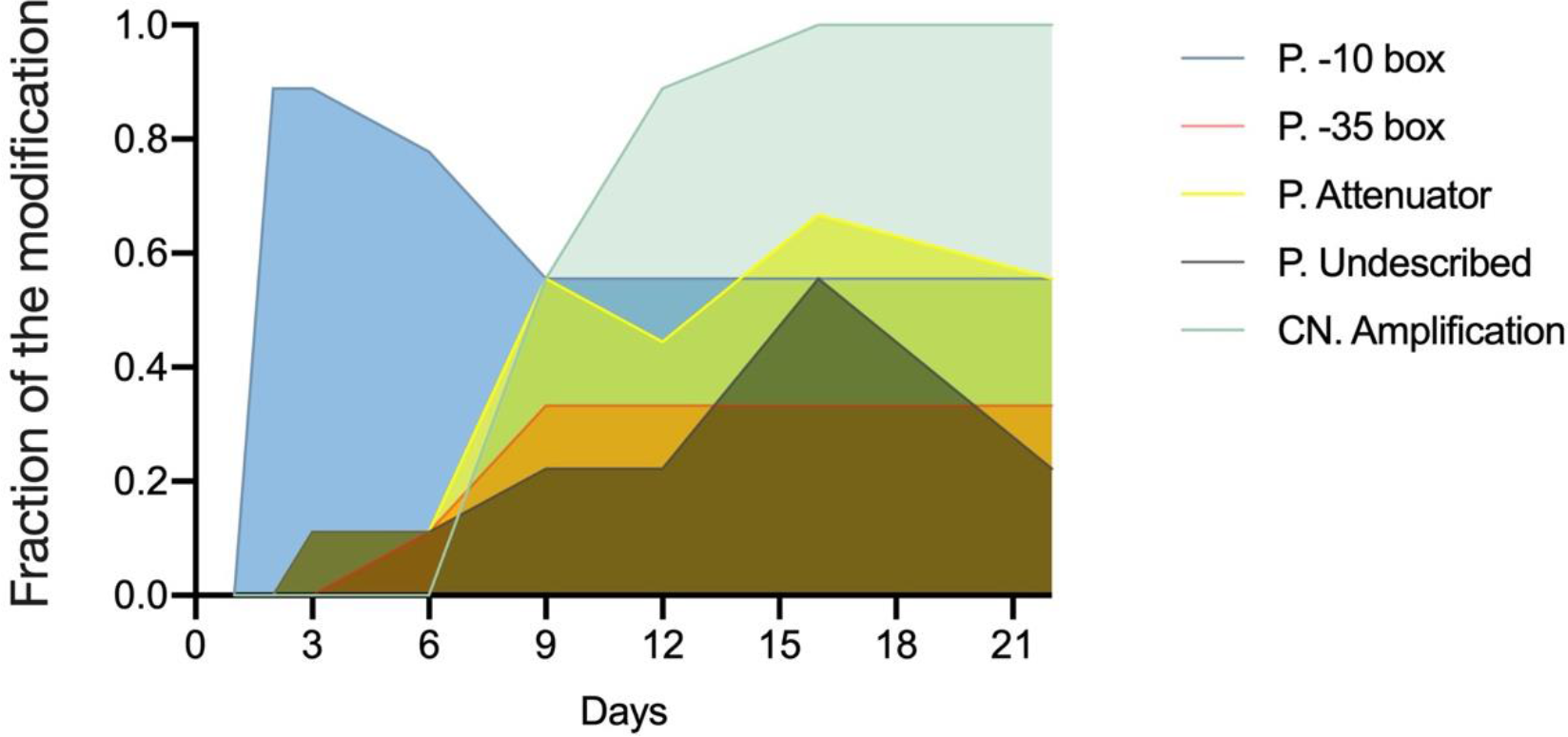
Trajectory of mutations in *ampC* promoter and copy number change. Mutations frequency was calculated in nine evolved population. P. represents *ampC* promoter. CN. represents *ampC* copy number. Undescribed is the mutations in other sites that have not been reported.

## DISCUSSION

Strains with higher AmpC production have a selective advantage in amoxicillin resistance development. The overexpression of the AmpC enzyme can be achieved by the enhancement of *ampC* gene transcription, both by increasing the *ampC* promoter strength and the gene copy number. When exposed to increasing concentrations of amoxicillin, *E. coli* gained resistance by increasing the transcription of *ampC* by a factor exceeding 100 (9). This study reports the time course of the amplification of a chromosomal fragment containing *ampC* and the genetic events accompanying it. The considerable delay before the first amplification events observed in this study, indicates that the mutated *ampC* promoter causes the initial increase of AmpC activity and amoxicillin resistance. Therefore, the process of amplification is not gradual. Instead, the initial jump in copy number seems to be around 8-fold and occurs after at least 6 days of exposure to increasing amoxicillin levels.

The mobile genetic elements could drive the antibiotic resistance genes’ copy number increase (15). The chromosomal DNA fragment isolated by Darphorn (16) from *E. coli* made amoxicillin resistant by de novo evolution, contains the IS1 transposon, which was introduced at the connection of the start and end of the fragments. This fragment can transfer from the amoxicillin evolved *E. coli* functioning as donor to susceptible *E. coli* receptor cells. In *Proteus mirabilia* amplification involving IS1 is based on homology recombination identifying two IS1 copies as homology regions for initial recombination, followed by tandem duplication of the region between IS1 elements (34). In this IS1-based amplification, the frequency of the initial duplication is 150-fold lower than that of the following amplification to higher copy numbers (35). If this is the same in *E. coli*, that would explain the pattern of *ampC* amplification observed here. All replicates with IS1 inserted in the junction of two amplification fragments had one of their amplifications flanking within the *ecnB* reading frame or promoter. The extremities of the IS1 sequence in particular are crucial for cointegration (36). The *ecnB* promoter and the tail of IS1 share the same 7 bp sequence. This implies that P*ecnB* contains a region homologous to part of IS1, suggesting a potential function of this sequence as the required homologous region for the recombination event, recruiting IS1 transposon and leading to amplification.

The absence of amplification in original position of *ampC* in evolved Δ*ampC* strains indicates that none of the other genes in the fragment confer enough resistance advantage for its amplification. It also rules out the already unlikely possibility that the cell would be aware of the location of *ampC* in the genome and amplify the region around it when exposed to amoxicillin. Instead, a large region, containing more than 1000 genes, was duplicated or triplicated. Similar amplification was also observed in tetracycline evolved *E.coli* (37), indicating that the driving factor may be the presence of multi-drug resistance genes within the amplified region.

Gene expression involves the coordination of multiple dynamic events subject to multi-level regulation (38). The positive relationship between AmpC activity and the MIC, *ampC* copy number and mRNA levels suggest that according to hierarchical control analysis (39) the genetic control component fully determines levels of expression, which in turn control AmpC activity. Hence, the *ampC* copy number exerts considerable influence over its expression, but that it is not the only factor. The mutations in the promoter areas are crucial in the initial stages of resistance development.

The comparison of gene mutations during evolution in WT and Δ*ampC* suggests that the deletion of *ampC* not only reduces the ability of resistance acquisition but also alters the evolutionary trajectory in *E. coli*. Point mutations in several genes exhibited higher frequency in evolved Δ*ampC* strains compared with WT, suggesting that additional mutations were needed to compensate for the missing *ampC* gene. The mutations in gene *ftsI* and *acrB* occur in all Δ*ampC* strains. These genes are known to confer amoxicillin resistance based on pathways associated with target alternation (40) and efflux pumps (30), respectively. In contrast, in the WT the mutation in *acrB* disappeared from the time point of the first amplification observed to the end of the evolution experiment in three replicates, and the mutation in *ftsI* was not observed. This suggests that these mutations may cause higher fitness costs than *ampC* amplification. Besides, mutations in *envZ* may confer resistance by decreasing drug influx through the membrane porins OmpC/OmpF (20)(41). Moreover, these mutations have been shown to increase carbapenem resistance in *ompCF*-deleted backgrounds (42). Mutations in the *envZ* gene occur in all evolved Δ*ampC* strains and also accumulate in several evolved WT strains, indicating that they synergize with AmpC.

Several DNA mutations present in the WT after resistance evolution were not observed in evolved Δ*ampC*. The mutations in *rpoD* accumulated in most evolved WT. However, this did not occur in evolved Δ*ampC*, indicating that *rpoD* may be important for *ampC* transcription enhancement. RNA polymerase coded by sigma factor 70 (σ^70^) gene *rpoD* is essential for gene transcription (43). RpoD connects to both the -10 and -35 regions in promoter to initiate transcription (44). A mutation at a same position in *rpoD* (Asp445Glu) was recently reported in cefotaxime evolved *E. coli* (32). Mutations in the *rpoD* gene in our study changed amino acids 445 (Asp445Ala, Asp445Val), 447 (Ala447Pro) and 570 (Asp570Gly) in several replicates. These sites are in the conserved regions 2.4 and 4.2 of the σ^70^ subunit, which connect to the -10 and -35 motif within the promoter, respectively (45). As mutations in gene *rpoD* occur later than *ampC* duplication and mutations in -10 box and -35 box of *ampC* promoter, the σ^70^ mutants potentially enhance binding affinity between the RNA polymerase and gene promoter and thereby improving the utilization of high gene dosage.

Altogether, our study can bring new thought of amoxicillin resistance prevention based on resistance emergence timeline. Also, we propose the DNA mutations which accompany with the *ampC* overexpression.

## MATERIALS AND METHODS

### Strains, culture conditions and adaptive evolution

Evans medium pH 6.9 supplemented with 55mM glucose (46) was used to culture the *E. coli* MG1655 (WT) and BW25113-Δ*ampC* (Δ*ampC*, NBRP *E. coli*, Keio library. Kanamycin resistance was removed) strains culturing at 37 °C. Stock solutions of amoxicillin (10 mg/mL) were dissolved with MM medium and filter-sterilized and stored at -80 °C. The evolved population everyday was stored with 30 % glycerol in -80 °C.

Wild-type *E. coli* Δ*ampC* were made amoxicillin evolved through laboratory evolution experiments as described before(47). Nine replicates of WT and three of Δ*ampC* were used for amoxicillin resistance evolution, and one culture not exposed to antibiotics was used as control. The starting amoxicillin concentration was 1 µg/mL for both WT and Δ*ampC*, which is a quarter of the WT *E. coli* MIC (4 µg/mL) and half of Δ*ampC* MIC (2 µg/mL), and the starting OD_600_ was 0.1. After 24 h grown in 10 mL tube with 5 mL Evans medium in 37 °C, 200 rpm incubator, if the OD_600_ of the culture in higher concentration was higher than 70 % that in lower concentration, the antibiotic concentration was doubled in the subsequent incubation, otherwise using the same antibiotic concentration. The evolution experiments lasted 22 days.

### MIC measuring

During the evolution process, MIC values were measured every 3 to 4 days in 96-well plates in plate readers (Thermo Scientific Multiskan FC with SkanIt software) according to previous description (48). Amoxicillin concentration ranged from 2 to 2048 µg/mL increasing by a factor of 2 at each step. The starting OD_600_ was 0.05. Plates were incubated at 37 °C for 24 h, with shaking and OD_595_ measurements were conducted every 10 min. The MIC was defined as the lowest amoxicillin concentration that reduced the growth to OD_595_ less than 0.2 after 24 h.

### AmpC activity measurement

The stored strains were grown in Evans medium overnight from storage tubes kept at -80 °C and diluted 1:100 with fresh Evans medium. Cultures were harvested at late-log phase and washed in 1 M PBS 7.0 buffer. Cells were diluted to OD_600_ 2 with PBS. 50 µL cell suspension was mixed with 50 µL 10 µg/mL nitrocefin and incubated at 37 °C in a plate reader with pulse shaking. The OD_492_ was measured every minute for 10 h. The fold-change of the activity was calculated by dividing the activity of evolved population by that of naive WT.

### Quantitative PCR

Genomic DNA was extracted with the DNeasy Blood and Tissue kit (Qiagen) for copy number measurements and whole genome sequencing. RNA was isolated using RNeasy Protect Bacteria Kit (Qiagen) and reverse transcription was carried out with iScript cDNA Synthesis kit (BIO-RAD).

TaqMan™ Universal PCR Mix (ThermoFisher) was used for Quantitative PCR (qPCR), performed with the Applied Biosystems 7300 realtime PCR system (Applied Biosystems) Primers and probes for qPCR (Table S5) were obtained from Integrated DNA Technologies and 6-FAM and TAMRA were used as dye and quencher of probe. A sample of naive WT was used as reference. The cDNA or genomic DNA were diluted to the same concentration (10 ng/µL). Cycle threshold (Ct) values were determined by automated threshold analysis using the ABI Prism 1.0 software. Gene copy numbers or gene relative production were determined using the -ΔΔCt method using GADPH as the reference gene.

### Sequencing of the *ampC* promoter

AmpC promoter was amplified by Herculase II Fusion DNA Polymerase (Agilent) using isolated genomic DNA as template and F-AmpC Prom, R-AmpC prom (Table S5) as primers. PCR product was purified using MSBSpinRapace kit (Stratec) and sequenced by Sangar (Macrogen Europe). The result was analyzed through Snapgene. Only the highest signal of mutations at each site was recorded.

### Whole-genome sequencing

Whole genome sequencing was conducted utilizing next generation sequencing Illumina (NextSeq 550 system) following established protocol(47). NEBNext Ultra II FS DNA Library Prep Kit for Illumina (New England BioLabs) and NEBNext Multiplex Oligos for Illumina (96 Unique Dual Index Primer Pairs; New England BioLabs) were used for creating a genomic DNA library. After removing the adapter using Cutadapt (49), the raw data was trimmed (50) and deduplicated. Then the bam files were aligned to references (NC000913 for WT and CP009273 for Δ*ampC*) with Bowtie2 (51). Freebayes (52) and Lofreq (53) were used for allele frequency calculation and variant calling. Snpeff (54)was used for variant annotation. The point mutations with allele frequency lower than 0.1 and those also occurring in the drug-free cultured control were removed.

The structure variation was confirmed through Breseq (55) and cn.mops (56). The trimmed fastq file was used for Breseq for transposition identification. The range of amplification fragment was determined from the output of cn.mops. These functions were conducted with bam file without deduplication according to Zhou et al (57).

## ACKNOWLEDGMENTS

The research was funded by the Netherlands Food and Consumer Product Safety Authority (NVWA). Luyuan Nong acknowledges the support from the Guangzhou Elite Project Scholarship. We thank Stanley Brul for his helpful suggestions. We also thank Selina Leeuwen and Alphonse de Koster for their assistance.

## REFERENCES

1. Ling LL, Schneider T, Peoples AJ, Spoering AL, Engels I, Conlon BP, Mueller A, Schäberle TF, Hughes DE, Epstein S, Jones M, Lazarides L, Steadman VA, Cohen DR, Felix CR, Fetterman KA, Millett WP, Nitti AG, Zullo AM, Chen C, Lewis K. 2015. A new antibiotic kills pathogens without detectable resistance. Nature 517:455–459.

2. Luong HX, Thanh TT, Tran TH. 2020. Antimicrobial peptides – Advances in development of therapeutic applications. Life Sci 260:118407.

3. Klein EY, Boeckel TPV, Martinez EM, Pant S, Gandra S, Levin SA, Goossens H, Laxminarayan R. 2018. Global increase and geographic convergence in antibiotic consumption between 2000 and 2015. Proc Natl Acad Sci 115:E3463–E3470.

4. Cantón R, Morosini M. 2011. Emergence and spread of antibiotic resistance following exposure to antibiotics. FEMS Microbiol Rev 35:977–991.

5. Dionisio F, Baquero F, Fuertes M. 2023. Psychological and cultural factors influencing antibiotic prescription. Trends Microbiol 31:559–570.

6. Jacoby GA. 2009. AmpC β-Lactamases. Clin Microbiol Rev 22:161–182.

7. Shaikh S, Fatima J, Shakil S, Rizvi SMohdD, Kamal MA. 2015. Antibiotic resistance and extended spectrum beta-lactamases: Types, epidemiology and treatment. Saudi J Biol Sci 22:90–101.

8. Honoré N, Nicolas MH, Cole ST. 1986. Inducible cephalosporinase production in clinical isolates of Enterobacter cloacae is controlled by a regulatory gene that has been deleted from *Escherichia coli*. EMBO J 5:3709–3714.

9. Händel N, Schuurmans JM, Brul S, Kuile BH ter. 2013. Compensation of the Metabolic Costs of Antibiotic Resistance by Physiological Adaptation in *Escherichia coli*. Antimicrob Agents Ch 57:3752–3762.

10. Jaurin B, Grundström T, Normark S. 1982. Sequence elements determining *ampC* promoter strength in *E. coli*. Embo J 1:875–881.

11. Andersson DI, Hughes D. 2009. Gene Amplification and Adaptive Evolution in Bacteria. Annu Rev Genet 43:167–195.

12. Normark S, Edlund T, Grundström T, Bergström S, Wolf-Watz H. 1977. *Escherichia coli* K-12 Mutants Hyperproducing Chromosomal Beta-Lactamase by Gene Repetitions. J Bacteriol 132:912–922.

13. Edlund T, Grundström T, Normark S. 1979. Isolation and characterization of DNA repetitions carrying the chromosomal β-lactamase gene of *Escherichia coli* K-12. Mol Gen Genetics Mgg 173:115–125.

14. Edlund T, Normark S. 1981. Recombination between short DNA homologies causes tandem duplication. Nature 292:269–271.

15. Maddamsetti R, Yao Y, Wang T, Gao J, Huang VT, Hamrick GS, Son H-I, You L. 2024. Duplicated antibiotic resistance genes reveal ongoing selection and horizontal gene transfer in bacteria. Nat Commun 15:1449.

16. Darphorn TS, Hu Y, Sintanneland BBK, Brul S, Kuile BH ter. 2021. Multiplication of *ampC* upon Exposure to a Beta-Lactam Antibiotic Results in a Transferable Transposon in *Escherichia coli*. Int J Mol Sci 22:9230.

17. Apostolakos I, Feudi C, Eichhorn I, Palmieri N, Fasolato L, Schwarz S, Piccirillo A. 2020. High-resolution characterisation of ESBL/pAmpC-producing *Escherichia coli* isolated from the broiler production pyramid. Sci Rep 10:11123.

18. Darby EM, Trampari E, Siasat P, Gaya MS, Alav I, Webber MA, Blair JMA. 2022. Molecular mechanisms of antibiotic resistance revisited. Nat Rev Microbiol 1–16.

19. Chowdhury N, Suhani S, Purkaystha A, Begum MK, Raihan T, Alam MdJ, Islam K, Azad AK. 2019. Identification of AcrAB-TolC Efflux Pump Genes and Detection of Mutation in Efflux Repressor AcrR from Omeprazole Responsive Multidrug-Resistant *Escherichia coli* Isolates Causing Urinary Tract Infections. Microbiol Insights 12:1178636119889629.

20. Choi U, Lee C-R. 2019. Distinct Roles of Outer Membrane Porins in Antibiotic Resistance and Membrane Integrity in *Escherichia coli*. Front Microbiol 10:953.

21. Lopatkin AJ, Bening SC, Manson AL, Stokes JM, Kohanski MA, Badran AH, Earl AM, Cheney NJ, Yang JH, Collins JJ. 2021. Clinically relevant mutations in core metabolic genes confer antibiotic resistance. Science 371.

22. Pinheiro F, Warsi O, Andersson DI, Lässig M. 2021. Metabolic fitness landscapes predict the evolution of antibiotic resistance. Nat Ecol Evol 5:677–687.

23. He G-X, Zhang C, Crow RR, Thorpe C, Chen H, Kumar S, Tsuchiya T, Varela MF. 2011. SugE, a New Member of the SMR Family of Transporters, Contributes to Antimicrobial Resistance in *Enterobacter cloacae*. Antimicrob Agents Chemother 55:3954–3957.

24. Bishop RE. 2000. The bacterial lipocalins. Biochim Biophys Acta (BBA) - Protein Struct Mol Enzym 1482:73–83.

25. Gottesman S. 2017. Stress Reduction, Bacterial Style. J Bacteriol 199.

26. Masi M, Pinet E, Pagès J-M. 2020. Complex Response of the CpxAR Two-Component System to β-Lactams on Antibiotic Resistance and Envelope Homeostasis in *Enterobacteriaceae*. Antimicrob Agents Chemother 64.

27. Kenney LJ, Anand GS. 2020. EnvZ/OmpR Two-Component Signaling: An Archetype System That Can Function Noncanonically. EcoSal Plus 9.

28. Nestorovich EM, Danelon C, Winterhalter M, Bezrukov SM. 2002. Designed to penetrate: Time-resolved interaction of single antibiotic molecules with bacterial pores. Proc Natl Acad Sci 99:9789–9794.

29. Misra R, Benson SA. 1988. Isolation and characterization of OmpC porin mutants with altered pore properties. J Bacteriol 170:528–533.

30. Kobayashi N, Tamura N, Veen HW van, Yamaguchi A, Murakami S. 2014. β-Lactam Selectivity of Multidrug Transporters AcrB and AcrD Resides in the Proximal Binding Pocket*. J Biol Chem 289:10680–10690.

31. Curtis NAC, Eisenstadt RL, Turner KA, White AJ. 1985. Inhibition of penicillin-binding protein 3 of *Escherichia coli* K-12. Effects upon growth, viability and outer membrane barrier function. J Antimicrob Chemother 16:287–296.

32. Schenk MF, Zwart MP, Hwang S, Ruelens P, Severing E, Krug J, Visser JAGM de. 2022. Population size mediates the contribution of high-rate and large-benefit mutations to parallel evolution. Nat Ecol Evol 6:439–447.

33. Kobayashi K, Tsukagoshi N, Aono R. 2001. Suppression of Hypersensitivity of *Escherichia coli acrB* Mutant to Organic Solvents by Integrational Activation of the acrEF Operon with the IS 1 or IS 2 Element. J Bacteriol 183:2646–2653.

34. Peterson BC, Rownd RH. 1983. Homologous sequences other than insertion elements can serve as recombination sites in plasmid drug resistance gene amplification. J Bacteriol 156:177–185.

35. Peterson BC, Rownd RH. 1985. Drug resistance gene amplification of plasmid NR1 derivatives with various amounts of resistance determinant DNA. J Bacteriol 161:1042–1048.

36. Machida Y, Machida C, Ohtsubo H, Ohtsubo E. 1982. Factors determining frequency of plasmid cointegration mediated by insertion sequence IS1. Proc Natl Acad Sci 79:277–281.

37. Qi W, Jonker MJ, Teichmann L, Wortel M, Kuile BHT. 2023. The influence of oxygen and oxidative stress on de novo acquisition of antibiotic resistance in *E. coli* and *Lactobacillus lactis.* BMC Microbiol 23:279.

38. Mitsis T, Efthimiadou A, Bacopoulou F, Vlachakis D, Chrousos G, Eliopoulos E. 2020. Transcription factors and evolution: An integral part of gene expression (Review). World Acad Sci J.

39. Kuile BH ter, Westerhoff HV. 2001. Transcriptome meets metabolome: hierarchical and metabolic regulation of the glycolytic pathway. FEBS Lett 500:169–171.

40. Bellini D, Koekemoer L, Newman H, Dowson CG. 2019. Novel and Improved Crystal Structures of *H. influenzae, E. coli* and *P. aeruginosa* Penicillin-Binding Protein 3 (PBP3) and N. gonorrhoeae PBP2: Toward a Better Understanding of β-Lactam Target-Mediated Resistance. J Mol Biol 431:3501–3519.

41. Waukau J, Forst S. 1992. Molecular analysis of the signaling pathway between EnvZ and OmpR in *Escherichia coli*. J Bacteriol 174:1522–1527.

42. Adler M, Anjum M, Andersson DI, Sandegren L. 2016. Combinations of mutations in *envZ, ftsI, mrdA, acrB and acrR* can cause high-level carbapenem resistance in *Escherichia coli*. J Antimicrob Chemother 71:1188–1198.

43. Helmann JD, Chamberlin MJ. 1988. Structure and Function of Bacterial Sigma Factors. Annu Rev Biochem 57:839–872.

44. Dombroski AJ, Johnson BD, Lonetto M, Gross CA. 1996. The sigma subunit of *Escherichia coli* RNA polymerase senses promoter spacing. Proc Natl Acad Sci 93:8858– 8862.

45. Barne KA, Bown JA, Busby SJW, Minchin SD. 1997. Region 2.5 of the *Escherichia coli* RNA polymerase σ^70^ subunit is responsible for the recognition of the ‘extended −10’ motif at promoters. EMBO J 16:4034–4040.

46. Evans CGT, Herbert D, Tempest DW. 1970. Chapter XIII The Continuous Cultivation of Micro-organisms 2. Construction of a Chemostat. Method Microbiol 2:277–327.

47. Qi W, Jonker MJ, Leeuw W de, Brul S, Kuile BH ter. 2023. Reactive oxygen species accelerate de novo acquisition of antibiotic resistance in *E. coli*. iScience 26:108373.

48. Schuurmans JM, Hayali ASN, Koenders BB, Kuile BH ter. 2009. Variations in MIC value caused by differences in experimental protocol. J Microbiol Meth 79:44–47.

49. Kechin A, Boyarskikh U, Kel A, Filipenko M. 2017. cutPrimers: A New Tool for Accurate Cutting of Primers from Reads of Targeted Next Generation Sequencing. J Comput Biol 24:1138–1143.

50. Bolger AM, Lohse M, Usadel B. 2014. Trimmomatic: a flexible trimmer for Illumina sequence data. Bioinformatics 30:2114–2120.

51. Langmead B, Salzberg SL. 2012. Fast gapped-read alignment with Bowtie 2. Nat Methods 9:357–359.

52. Garrison E, Marth G. 2012. Haplotype-based variant detection from short-read sequencing. arXiv.

53. Wilm A, Aw PPK, Bertrand D, Yeo GHT, Ong SH, Wong CH, Khor CC, Petric R, Hibberd ML, Nagarajan N. 2012. LoFreq: a sequence-quality aware, ultra-sensitive variant caller for uncovering cell-population heterogeneity from high-throughput sequencing datasets. Nucleic Acids Res 40:11189–11201.

54. Cingolani P, Platts A, Wang LL, Coon M, Nguyen T, Wang L, Land SJ, Lu X, Ruden DM. 2012. A program for annotating and predicting the effects of single nucleotide polymorphisms, SnpEff. Fly 6:80–92.

55. Gifford DR, Furió V, Papkou A, Vogwill T, Oliver A, MacLean RC. 2018. Identifying and exploiting genes that potentiate the evolution of antibiotic resistance. Nat Ecol Evol 2:1033–1039.

56. Klambauer G, Schwarzbauer K, Mayr A, Clevert D-A, Mitterecker A, Bodenhofer U, Hochreiter S. 2012. cn.MOPS: mixture of Poissons for discovering copy number variations in next-generation sequencing data with a low false discovery rate. Nucleic Acids Res 40:e69– e69.

57. Zhou W, Chen T, Zhao H, Eterovic AK, Meric-Bernstam F, Mills GB, Chen K. 2014. Bias from removing read duplication in ultra-deep sequencing experiments. Bioinformatics 30:1073–1080.

